# Two-step multi-omics modelling of drug sensitivity in cancer cell lines to identify driving mechanisms

**DOI:** 10.1101/2020.08.28.271544

**Authors:** Nina Kusch, Andreas Schuppert

## Abstract

Drug sensitivity prediction models for human cancer cell lines constitute important tools in identifying potential driving factors of responsiveness in a pre-clinical setting. Integrating information derived from a range of heterogeneous data is crucial, but remains non-trivial, as differences in data structures may hinder fitting algorithms from assigning adequate weights to complementary information that is contained in distinct omics data. In order to counteract this effect that tends to lead to just one data type dominating supposedly multi-omics models, we developed a novel tool that enables users to train single-omics models separately in a first step and to integrate them into a multi-omics model in a second step. Extensive ablation studies are performed in order to facilitate an in-depth evaluation of the respective contributions of singular data types and of combinations thereof, effectively identifying redundancies and interdependencies between them. Moreover, the integration of the single-omics models is realized by a range of distinct classification algorithms, thus allowing for a performance comparison. Sets of molecular events and tissue types found to be related to significant shifts in drug sensitivity are returned to facilitate a comprehensive and straightforward analysis of potential drivers of drug responsiveness. Our two-step approach yields sets of actual multi-omics pan-cancer classification models that are highly predictive for a majority of drugs in the GDSC data base. In the context of targeted drugs with particular modes of action, its predictive performances compare favourably to those of classification models that incorporate multi-omics data in a simple one-step approach. Additionally, case studies demonstrate that it succeeds both in correctly identifying known key drivers of specific drug compounds as well as in providing sets of potential candidates for additional driving factors of drug sensitivity.

## Introduction

Large-scale pharmacogenomic cell line data bases featuring both in-depth multi-omics characterizations and extensive pharmacological profiles of human cancer cell lines constitute a crucial tool in uncovering potential driver mechanisms of drug sensitivity towards anti-cancer drug compounds [1–3]. To this end, multiple studies have been conducted building predictive models based on a range of different omics data types separately to predict both pre-clinical and clinical drug sensitivity, including baseline gene expression patterns [4, 5], somatic mutations [1], copy number variations (CNVs) and hypermethylation events [6, 7], or tissue lineage [2, 3].

Moreover, several comparative studies have been performed that assess the respective impact, benefits and shortcomings of specific modelling choices in the context of cancer drug sensitivity prediction tasks: Working with a set of human breast cancer cell lines, Costello et al. evaluate 44 drug sensitivity prediction algorithms proposed in the framework of a DREAM challenge, including among others kernel methods, non-linear, sparse linear and principal-component regression approaches, as well as ensemble models [8]. They find that leveraging all available omics data in addition to integrating external information, related for instance to biological pathways, improves prediction performance, as does employing non-linear modelling approaches. Gene expression data is found to constitute the most potent predictor variable, potentially as a consequence of its data structure and the wealth of customized tools available to process it. Jang et al. systematically assess the performance of distinct choices in five components of the modelling pipeline, including the choice of input features and the choice of fitting algorithm, as well as the overall impact of these components themselves on model predictivity [9]. Concluding that the most important modelling factor is the choice of features and agreeing with Costello et al. on the dominance of gene expression data, they rate the choice of algorithm as the third most important modelling factor. Expanding on this concept of systematically identifying optimal choices in distinct steps of the modelling pipeline and applying it to translative modelling, Turnhoff et al. have published an R package that can be used to perform even more intricate analyses in the context of predicting clinical responses while training on cell line data [10, 11].

In addition to the missing consensus on optimal fitting strategies, another unsolved problem is how to adequately integrate heterogeneous data types into one common model while assigning the proper weights to complementary information from distinct data. To this end, Aben et al. propose a two-stage approach that first utilizes upstream omics data and consequently fits the resulting errors with a second model based on gene expression [12, 13]. By employing this method on the GDSC data set, they aim to counteract the tendency of gene expression data to dominate models that are designed to integrate information from distinct omics data types in a straightforward approach [1, 2, 7–9, 14, 15]. This ansatz, however, fails to incorporate pathway information and impedes a simple quantitative analysis into the relative importance of the contributions of the entirety of data types to the model and consequently, into the relative influence of possible driving mechanisms that determine drug sensitivity by focusing mostly on the subset of upstream omics data. Their results demonstrate that switching from a one-stage to a two-stage modelling approach yields negligible changes in predictive performance, but can produce models that are more easily understandable, which is imperative for translating the results to a clinical setting.

In this paper, we present a two-step modelling approach to classify pre-clinical drug sensitivity in cancer based on six distinct feature types, namely basal gene expression, somatic mutation, CNV and hypermethylation events, pathway activation scores provided by the R package PROGENy [16, 17], and information on tissue lineage. A set of models is first trained on one data type each, before their respective outputs are collectively utilized as input features in a second step, where a range of different classification algorithms are employed, including a Naïve Bayes classifier, a shallow Neural Network, Support Vector Machines, decision tree ensembles, and both linear and logistic regression approaches with multiple distinct regularization schemes. This ansatz leverages and applies the full range of crucial insights gained from the aforementioned studies, such as the importance of integrating distinct and complementary data types, especially pathway information and gene expression data, and the challenge of realizing this when fitting models on all structurally heterogeneous data types simultaneously. In turn, our approach contributes substantially to the field of pre-clinical classification models of drug sensitivity in cancer: In addition to enabling the user to construct pan-cancer models that integrate various potentially complementary molecular and genetic data types in a way that impedes any type to overpower the others based solely on its structural properties, our ansatz also allows for a straightforward in-depth analysis of the respective contributions of data types and combinations of data types as well as the impact of singular features on drug sensitivity. Both the code of this tool as well as the data discussed in this paper are publicly available and can be downloaded at https://github.com/JRC-COMBINE/two-step-modelling.

## Materials and methods

### Data

The data to train and test the models on was originally generated in the context of the Genomics of Drug Sensitivity in Cancer project (GDSC) [1, 18] and has been partially processed by Iorio et al. [7, 19]. We use basal gene expression data, information on somatic mutation, CNV and hypermethylation events as well as tissue descriptors of cell lines as input features for our models. Moreover, we downloaded the PROGENy R package developed by Schubert et al. [16, 17] and applied it to the gene expression data in order to infer pathway activation scores. We obtained the area under the dose-response curve (AUC) values as measure for drug sensitivity and detailed annotations of genes, cell lines and drugs from the GDSC project. A detailed list of the files downloaded and an in-depth description of processing steps can be found in Table 1.

**Table 1.**
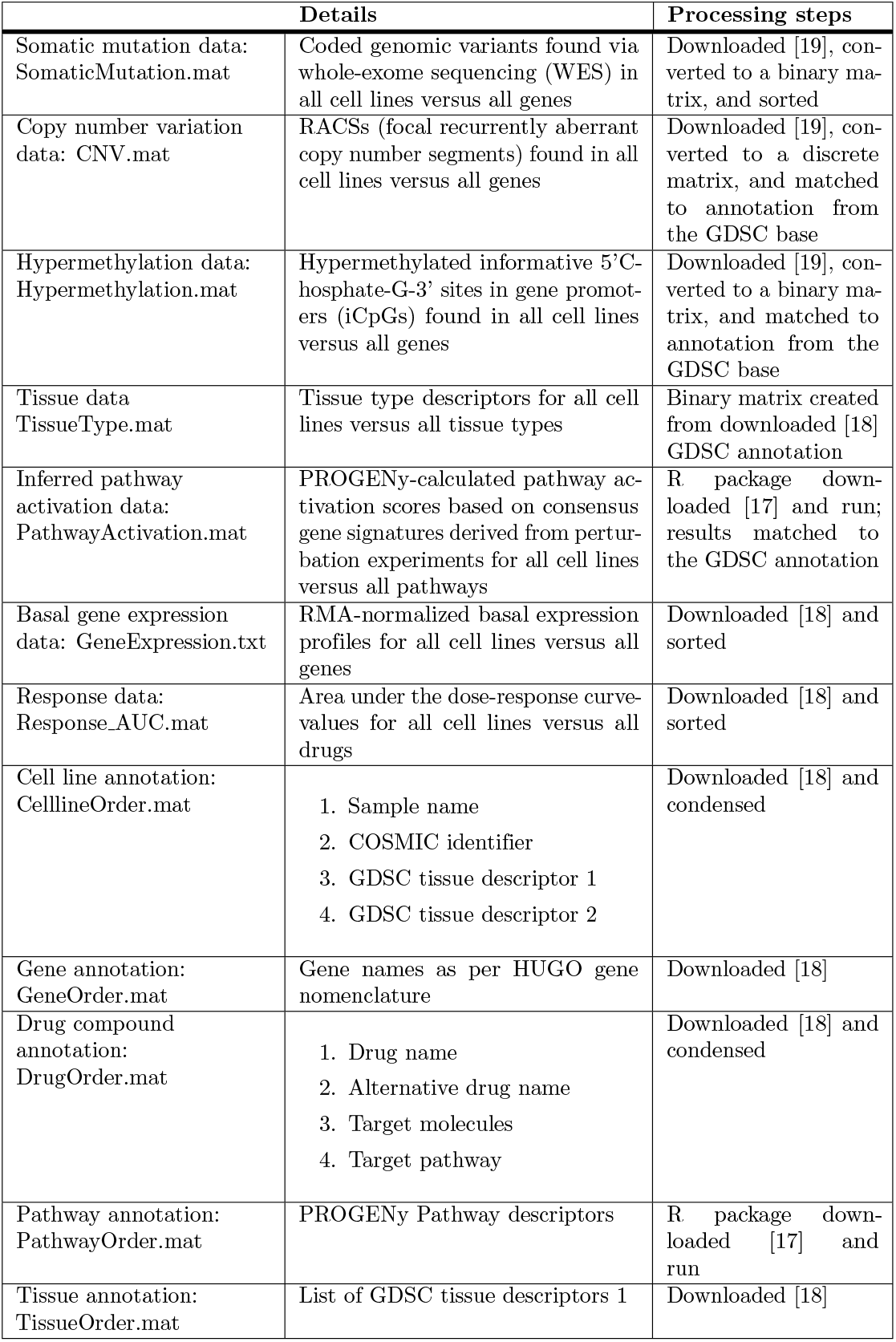
Data files used for model fitting and the respective processing

### Implementation

While Fig 1 provides a simplified overview of the model workflow, a more comprehensive visualization can be found in S1 Fig. The MATLAB routine twostepmodel.m is called with one input variable, namely the index of the drug compound to be modelled in the GDSC data base, an integer between 1 and 265. The drug compound annotation, as described in Table 1, holds the names of and additional pieces of information about the compounds corresponding to any such index.

**Fig. 1.**
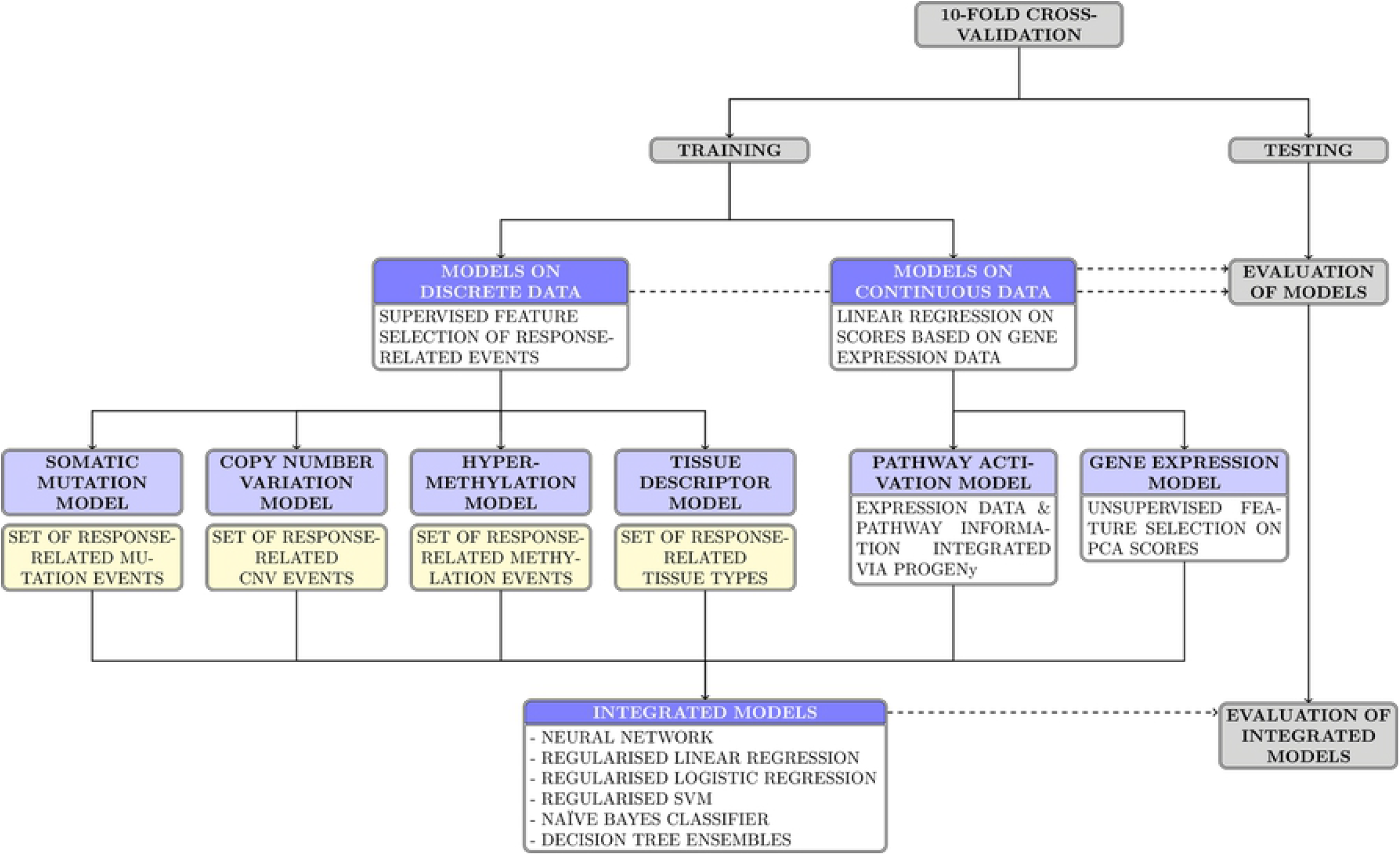
Two-step modelling workflow. A simplified diagram of the two-step modelling workflow, visualizing the link between the single-data type first-step models and the integrated second-step models. Boxes shaded in dark blue symbolize sets of models, while light-blue boxes represent individual models. Yellow boxes stand for sets of discrete features that are linked to a shift in drug sensitivity. Ablation models are not included in the graphic.

The two-step modelling routine computes three sets of drug-specific models across ten cross-validation folds: six first-step models that are based on one single data type each, 13 integrated second-step models that use all non-constant outputs of the first-step models as inputs, and up to 41 ablation models per second-step model. The latter result from applying all 13 fitting algorithms to reduced sets of inputs with up to three input vectors missing. Model performances are evaluated by a range of different metrics, including predictive accuracy and ROC-AUC, with the latter not being computed for models based on the Naïve Bayes classifier. Singular discrete features of interest that are found to be linked to a shift in drug sensitivity are returned to the user, as are the weights and importance scores for the outputs of the first-step models, as calculated by the second-step fitting algorithms. The twelve outputs of the routine are listed and explained in Table 2 and are designed to enable the user to comprehensively analyse the resulting models and their respective features and performance.

**Table 2.** Model outputs

| Quantity of interest | Details |
| --- | --- |
| First-step models | Predicted response for all clusters calculated by the discrete first-step models; pathway activation- and gene expression-based model; feature values associated with the genetic-feature based clusters |
| Second-step models | 13 integrated multi-omics models |
| Predictions on the training set | Binarized predictions of all first-step models and second-step models; pre-binarized predictions of all first-step models and all second-step models minus the Naïve Bayes model |
| Predictions on the test set | Binarized predictions of all first-step models and second-step models; pre-binarized predictions of all first-step models and all second-step models minus the Naïve Bayes model |
| Measured response | Measured response data, both unprocessed and binarized, in the training and in the testing set |
| Model performance on the training set | Evaluation metrics – accuracy, precision, recall, f1-score, FDR – for all first- and second step models in training |
| Model performance on the test set | Evaluation metrics – accuracy, precision, recall, f1-score, FDR – for all first- and second step models in testing |
| ROC-AUCs | ROC-AUCs of all first- and second-step models, with the exception of the Naïve Bayes classifier, in training and testing |
| Significant features | Sets of relevant mutation, CNV, and methylation events as well as tissue types used in the discrete first-step models, in addition to the three extended sets of genetic features, including redundant features |
| Importance of first-step models | Indices of all non-constant first-step models; input weights as calculated by the linear and logistic regression models and the SVMs as well as input importance scores calculated by the ensemble models |
| Ablation studies | Up to 41 ablation models for each second-step model; ROC-AUCs, if applicable, and accuracy values of all ablation models in training and testing |
| Cross-validation | Partition object used for the 10-fold cross validation |

Descriptions of outputs are listed in the order of them being returned by the routine; if not stated differently, all featured quantities are computed for each of the 10 cross-validation folds, which correspond to rows in the resulting output objects.

Any results pertaining to the first-step models are ordered as follows: somatic mutation-based, CNV-based, hypermethylation-based, tissue-based, pathway activation-based and gene expression-based. Results of second-step models are ordered in the same way that the corresponding algorithms are listed in the supplementary table S1 Table, which contains a detailed list of the 13 algorithms used as well as their parameter settings, whenever they diverge from the default settings provided by MATLAB.

#### First-step models

The first-step models that predict drug sensitivity based on one of the four discrete data types – somatic mutation, CNV, hypermethylation and tissue descriptors – are structurally similar, as are the data they are utilizing: Both the hypermethylation and the CNV matrix of cell lines versus genes are sparse, especially the latter, and they as well as the somatic mutation matrix include columns that are highly correlated and therefore contain information redundant for prediction purposes. The binary matrix holding information about the tissues to which the cell lines belong according to the descriptors used in the GDSC annotation features a significantly lower number of columns, all of which are linearly independent.

All discrete first-step models aim to identify features that are associated with a shift in responsiveness between cell lines that display such a feature versus those that lack it. To this end, all features that can be used to split the set of all training cell lines in two such subsets of sufficiently large size are tested for such an association. In the case of the three genomic feature data types, namely mutation, CNV, and hypermethylation data, all genes are screened to check if at least 15 instances of at least two distinct states are present in the training data; if not, they are temporarily discarded. For tissue data, tissue types are considered if they are featured at least ten times in the training set. A two-sided t-test with a significance level of *α* = 0.05 and Bonferroni-correction is conducted on the pre-binarization response data to identify significant differences in responsiveness between the sets of cell lines where a specific feature is either present or absent. All genes yielding statistically significant results are used to sort the training cell lines into clusters where all members exhibit the exact identical pattern of significant features being present or absent. If no such feature is identified, all training cell lines are pooled into one trivial cluster and their mean binarized responsiveness is set as the model prediction for all cell lines indiscriminately. Any model yielding such a constant prediction vector that is devoid of information is subsequently removed and not utilized as input to the second-step models.

Models identifying at least one feature significantly associated with a shift in responsiveness use these to sort all training cell lines into *2^n^* clusters, where *n* denotes the number of non-redundant features found. Pairs of redundant features, that is two features both present and absent in exactly the same subset of training cell lines, are identified, and one of them is removed from the set of relevant features used for clustering. Empty clusters are discarded, and for all remaining ones, the mean responsiveness over all training cell lines associated with a cluster is computed. Subsequently, it is used as the model prediction for any test cell line that would belong to the respective cluster based on their profile of relevant features. Should the somatic mutation-based model based identify two significant, non-redundant features, for instance mutations in TP53 and BRAF, it would sort all training cell lines into four clusters: the cluster of cell lines where both mutations are present, that of cell lines where both mutations are absent, and two clusters of cell lines where exactly one of these two mutations would be present.

In order to speed up computation times, an upper limit to the number of non-redundant relevant features has been introduced for mutation, CNV and hypermethylation data. The routine returns the set of relevant features as well as the reduced set of relevant, non-redundant features that is ultimately used for cluster definition and drug sensitivity prediction. In addition, the user can access the percentage of responding cell lines associated with each cluster as well as the exact state of any relevant, non-redundant feature in each cluster.

In the case of tissue data, redundancy cannot occur and cell lines are sorted into *n* +1 clusters if *n* tissue types are found to be associated with a significant change in responsiveness: the last *n* clusters correspond to the *n* tissue types and the first one is composed of the cell lines of all remaining tissues. This ordering applies to all outputs pertaining to the clusters of the tissue models: for instance, the average percentage of responding cell lines per cluster starts with that of the joined cluster of non-significant tissue types and continues with the values corresponding to the singular-tissue clusters in the order that these tissues are listed in the sets of significant features. A schematic visualization of the structure of the first-step models that utilize the discrete data types can be found in Fig 2.

**Fig. 2.**
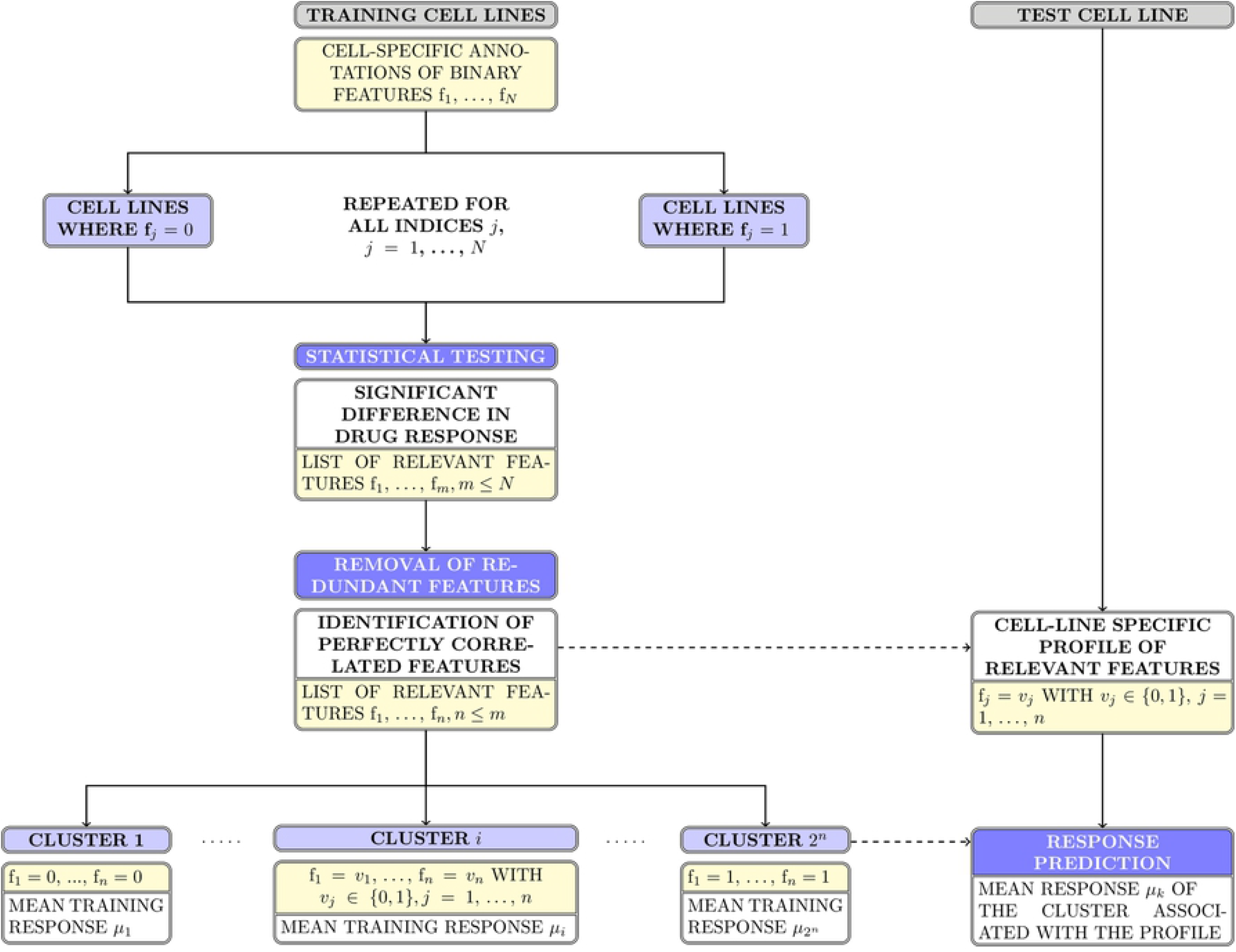
First-step models on discrete data types. Schematic visualization of the structure of the discrete first-step models built on genomic features. In the case of the CNV-based model, single features f_*j*_ may also take on values in {0, −1}, with the latter value denoting a deletion event. The tissue-based model is structured similarly, but does not require the removal of redundant features and defines *n* +1 clusters out of *n* features identified as significant. Dotted lines represent routines that are computed on the training data and subsequently applied to the test data set. Boxes in yellow refer to sets of features, while those coloured light blue indicate subsets of cell lines.

As for the two continuous molecular data types, a principal component analysis is performed on the basal gene expression data of the training cell lines and the first seven components are used to fit a linear regression model to the binarized response data. The number of principal components was chosen based on an analysis of their respective variances, as computed on the complete data set; in particular, it was determined to constitute a fitting trade-off between including the highest possible number of potentially informative components and reducing the number of input vectors in order to facilitate the fitting process. A second linear regression model is fitted on the eleven pathway activation scores provided by the PROGENy package for the training cell lines. The output vectors of these two models are normalized to the interval [0, 1] and an optimal binarization cutoff is calculated such that the predictive accuracy is maximized.

#### Second-step models

After the computation of the six first-step models, constant model output vectors – the results of models that failed to find any statistically significant relation between drug sensitivity and at least one genomic event or tissue – are removed, while the remaining ones are used as input feature vectors for 13 fitting algorithms in a second step. These algorithms comprise both one straight-forward classification approach, namely a Naïve Bayes classifier, as well as a diverse set of regression methods that are post-processed to identify an optimal cut-off yielding a classification with maximal accuracy. For these models, an additional performance evaluation metric is computed, the ROC-AUC, as detailed in Table 1.

For a subset of second-step models, weights and importance scores can be calculated in order to estimate the significance of the contributions of singular data types; in those cases, namely all algorithms but the Neural Networks and the Naïve Bayes classifier, these coefficients are returned to the user. Since at least three of the data types used have been shown to be interdependent and to contain highly redundant information, these fitted coefficients may not accurately reflect the actual significance of the information contained in any particular data type. In order to address this effect and to gain an improved insight into this issue, ablation studies are performed subsequently.

#### Ablation studies

Since tissue types and genome-wide expression patterns such as principal components have been shown to coincide [20] and since the PROGENy-derived pathway activation scores are calculated by integrating pathway information and gene expression, we expect these data types to contain highly correlated and redundant information. As a consequence, ablation studies are performed where all possible combinations of up to three first-step model outputs are omitted as inputs to the second-step fitting algorithms, which are then run on the remaining inputs. In the case of all six data types producing non-constant first-step models, a set of 41 ablation models is computed for any of the 13 second-step fitting algorithms. Accuracy and ROC-AUC values are calculated for any ablation model except for those applying a Naïve Bayes classifier, where only the accuracy is calculated.

## Results and discussion

Running the proposed two-step modelling routine on any of the 265 drug compounds present in the GDSC data set results in 13 two-step integrated multi-omics models, six first-step models based on one data type only, and up to 41 ablation models with combinations of up to three first-step models removed for any of the aforementioned 13 algorithms. Additionally, sets of events associated with shifts in responsiveness, performance evaluation metrics in training and testing and the weights or importance associated with first-step models by a subset of the second-step models are computed and returned to the user. All of these results are calculated for each of the 10 folds used in the cross-validation scheme that is applied to the data. A detailed list of the entirety of output objects and details about the exact pieces of information they hold is presented in Table 2.

When computing the averaged test ROC-AUC over the cross-validation folds of each model calculated on the 265 drug compounds, with the exception of the Naïve Bayes classifier, we achieve a spread ranging from 0.92 for the best model across all drugs to 0.37 for the model yielding the overall worst test performance. The worst performance over all drug-specific best models results in a mean test ROC-AUC of 0.50, while for 155 drugs, at least one model produces a mean test ROC-AUC of at least 0.7. A visualization of these findings is provided in Fig1–3 in S1 Appendix; for any particular drug compound, the distribution of the ten test ROC-AUCs of the one model yielding the highest mean test ROC-AUC is shown.

**Fig. 3.**
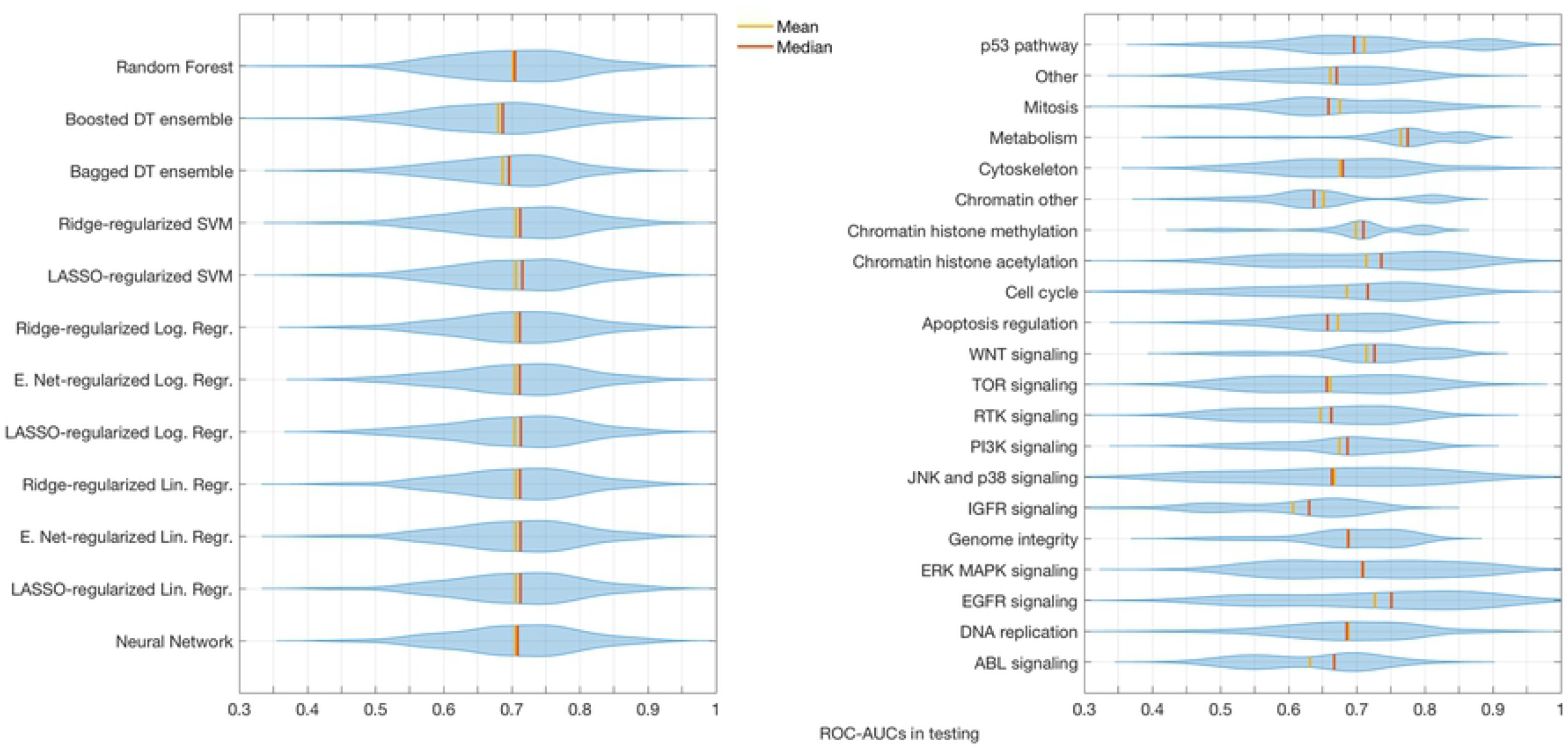
Effects of the choice of fitting algorithms and target drug classes on predictive performances. Distribution of mean test ROC-AUCs of models using different fitting algorithms for the second-step models (left) and of models being fitted to drug compounds of distinct classes, annotated by the target structure (right).

### Impact of drug classes and algorithms on the model performances

Evaluating the distribution of the averaged test ROC-AUCs over all drug compounds for all suitable two-step models separately, we observe no significant difference between the distinct second-step fitting algorithms when correcting for multiple testing. Notable are slightly lower means and medians for both of the decision tree ensembles and a lack of models producing mean test ROC-AUCs over 0.87 for the bagged decision tree ensembles. In addition, we find that the boosted decision tree ensembles exhibit signs of severe overfitting in drugs that feature less than 500 measurements across cell lines, as depicted in Fig 1 in S2 Appendix. When excluding models fitted via boosted decision tree ensembles, the remaining models tend to exhibit gradually fewer cases of overfitting as their averaged training ROC-AUC increases, an effect that is visualized in Fig 2 in S2 Appendix.

In contrast to the choice of second-step algorithm, the type of drug compound to be modelled greatly influences the performance of the resulting models. Using the official annotation, the drug compounds featured in the GDSC data base can be sorted into 20 well-defined classes with respect to their target pathway or mechanism and one more broadly-defined class titled ‘other’. After binning together all models calculated on any drug compound of a particular class, a two-sided t-test with a Bonferroni-correction for multiple testing is applied to the respective mean test ROC-AUC values and yields a high number of significantly different results between drug classes. These findings replicate the conclusions drawn by Jang et al [9] upon working with a prior version of the GDSC data set, namely that the choice of input features and the compound to be modelled exert a far stronger influence on the variance of predictive accuracy than the choice of algorithm. The results of a statistical analysis of this effect can be found in S3 Appendix, while a visualization of the variation in model performance across different algorithms and drug classes can be drawn from Fig 3.

### Comparison to one-step multi-omics classification models

The two-step multi-omics modelling approach presented in this paper is designed to create drug compound-specific classification models of cancer cell lines that integrate a range of distinct data types. This integration of heterogeneous data is realized in a separate second step that minimizes the chance that any inherent structural difference, such as sparsity or range, causes one data type to overpower additional ones and to drown out crucial information. As a consequence, complementary information contained in structurally heterogeneous data types can be conserved and utilized to not only improve the predictive power of the models, but also to provide insight into the drivers of responsiveness. Our routine additionally provides the opportunity to analyse and quantify the importance of data types and to identify redundancies between the information they contain in a straightforward and easily interpretable manner.

In order to contextualize the overall predictive performance of the resulting classification models, the test ROC-AUC of any model is compared to the results obtained by Jang et al. [9]. In their study, 114,000 classification models are computed and applied to prior versions of the CCLE and the GDSC data sets with the aim to systematically quantify the importance of five categories in the modelling workflow. To the best of our knowledge, these results constitute the most fitting standard to compare the performance of our algorithm to, since they, in contrast to other studies, fit multi-omics pan-cancer classification models on the GDSC data set, using similar sets of data types as predictors as well as AUC values as the metric quantifying drug sensitivity. As evidenced by Aben et al. [12], we do not necessarily expect the two-step modelling approach to yield significantly improved predictive performances, since the algorithm was designed mainly to produce actual multi-omics models that enable an intuitive and straightforward analysis of data contributions.

While our study uses a more current version of the GDSC data base with 265 instead of 138 drugs and a higher number of input data types that additionally include hypermethylation events, pathway activation scores and somatic mutation events identified on all of the 17,737 genes present in the data set, comparability between the studies is ensured by a number of steps. Firstly, a similar binarization regime on the response data is applied by using upper and lower quartile thresholds, which results in a number of cell lines to be modelled that is reasonably close to the one achieved by Jang et al., who worked with a smaller number of cell lines and used tertiles as thresholds. Secondly, for any model we calculate test ROC-AUCs by computing the concatenated prediction vector over all each cross-validation folds and by comparing it to the vector of measured responses that are binarized using the entire data set. In internal evaluations, we prefer to study test ROC-AUCs averaged across all cross-validation folds, as the model routine binarizes the measured response vectors by calculating the thresholds on the training data only. Binarizing on the complete data set tends to result in a small number of cell lines being labelled differently and consequently does not accurately reflect the model performance; however, we treat these small alterations as negligible in order to ensure a fair comparison. Lastly, we apply a cross-validation regime and utilize a significant set of algorithms overlapping with the ones used by Jang et al., such as regularized regressions, random forests and support vector machine approaches.

The findings of the Jang study demonstrate that the choice of genomic features used to build a model and the drug compound to be predicted explain by far the most significant share of variation in model performance across the complete set of models; the choice of the fitting algorithm is found to be only the third most important with a considerably smaller influence on the results. As evidenced by Fig 3 and S3 Appendix, our study confirms that different drug classes can be associated with significantly different distributions of performance measures, while there is little variation observed between distinct fitting algorithms.

The results obtained for 138 distinct drug compounds that are present in both versions of the GDSC data set and included in both studies are compared by selecting for each drug compound the model yielding the best test ROC-AUC. The two-step modelling approach produces higher-ranking performances for 23 drug compounds, with a statistically significant enrichment for drugs targeting the EGFR signalling pathway. The classes of drugs targeting the p53, the WNT, as well as the JNK and p38 signalling pathway additionally yield statistically significant enrichment values in a single-test setting. This hints at our two-step approach potentially being particularly useful and capable of improving on the benefits of already existing modelling platforms in applications involving targeted drugs with particular modes of action. A detailed visualization of these results can be found in Fig 4 and S4 Appendix.

**Fig. 4.**
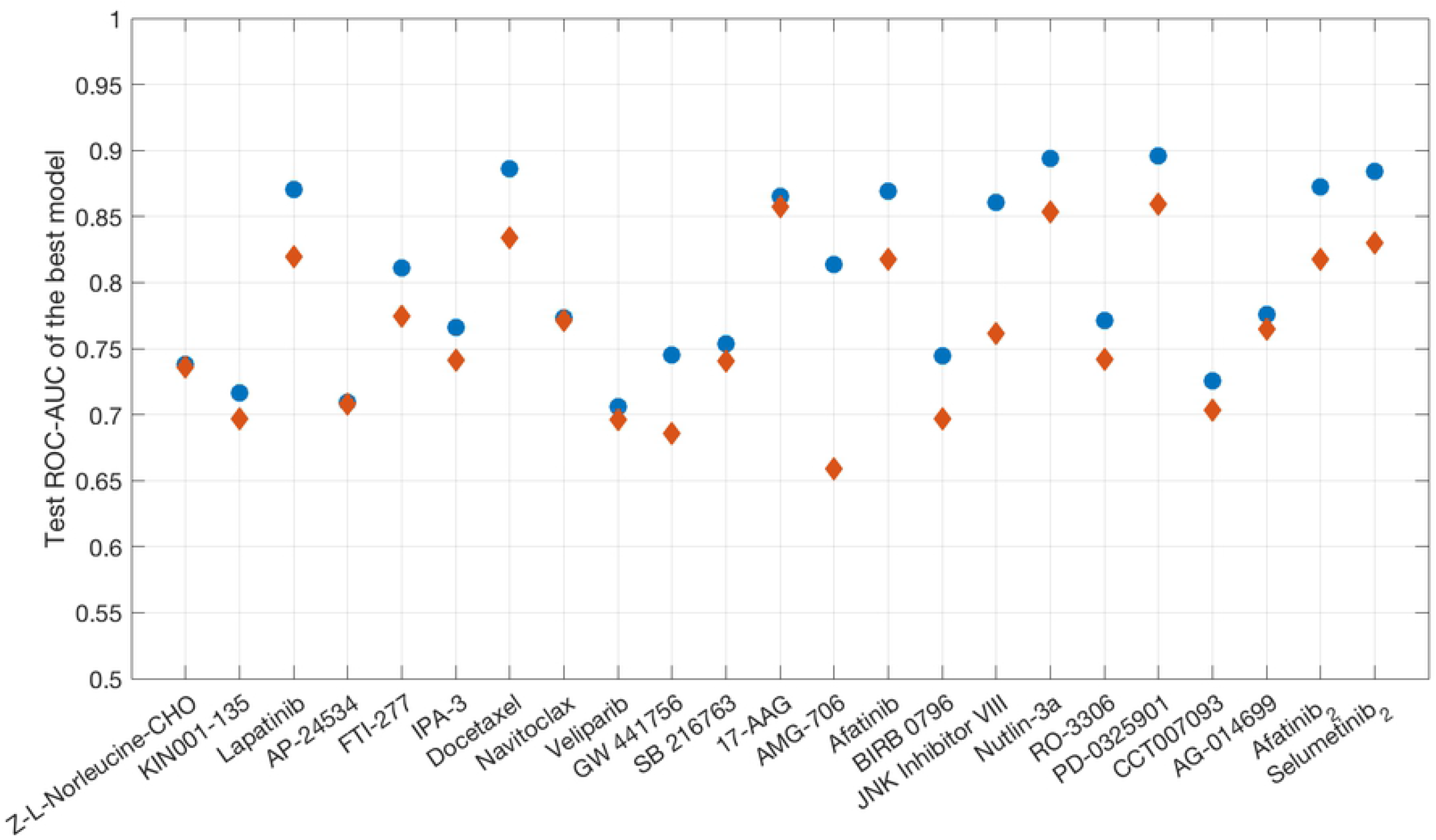
Two-step models outperforming one-step models. Concatenated test ROC-AUC of the respective best-performing two-step model (blue) for any drug where at least one two-step model outperforms all straightforward multi-omics models generated in the study of Jang et al. (red).

In contrast to the approaches applied by Jang et al, our two-step models additionally allow for an in-depth analysis of the absolute and relative contributions of data types – the weightiest component of the modelling workflow – to the resulting model performance and yield a list of the specific features that drive resistance or sensitivity to a particular drug. Not only can the performance of any two-step model integrating different data types be directly compared to that of up to six first-step models built on one data type only in order to quantify the benefits of incorporating additional input data, but extensive ablation studies enable the user to further analyse potential drops in model performance upon excluding combinations of up to three input data types. Thus, redundant information that is featured in more than one data type may be identified and the contribution of that data type to model performance can be quantified more accurately. Moreover, weights and importance scores assigned to data types by eleven of the second-step fitting algorithms can be assessed and studied.

### Case study 1: Nutlin-3a

Nutlin-3a is a small cis-imidazoline molecule and, a as potent inhibitor of *MDM2-TP53* interactions, known to induce senescence in cancer cells that express wild-type *TP53* [21], therefore repressing tumour growth in the absence of mutations of the *TP53* gene. It has been studied in pre-clinical settings in the context of a wide range of cancer types [22, 23]. Due to its well-understood mode of action, it constitutes a highly suitable candidate for evaluating whether the two-step modelling algorithm succeeds in correctly identifying drivers of drug responsiveness and in leveraging them in order to predict drug efficacy. In particular, we expect the somatic mutation-based model to outperform all other first-step models and to identify a mutation in *TP53* as strongly linked to a shift in responsiveness to Nutlin-3a.

Overall, the two-step modelling approach yields integrated models producing high ROC-AUCs in testing, with the best-performing models, namely the logistic regression approach with ridge or Elastic Net regularization, achieving a median ROC-AUC of 0.9 across all folds. In contrast, the majority of first-step models based on a singular data type struggle to classify the cell lines correctly into responders and non-responders, with the CNV-based, the hypermethylation-based, the tissue-based and, to a lesser extent, the gene expression-based models producing median ROC-AUCs in the range of 0.54 to 0.67. The sole models yielding moderate to high median ROC-AUCs are the somatic mutation-based and the pathway activation-based models with values of 0.86 and 0.8, respectively, as visualized in Fig 1 in S5 Appendix. Consequently, the outputs of these two models are assigned the highest averaged weights and importance factors, when fed as input features into the second-step fitting algorithms, as evidenced by Tables 1–2 in S5 Appendix. The remaining model outputs receive notably lower scores, as is consistent with their inferior predictive performance. A notable exception is the tissue-based model, which is assigned the lowest averaged weight, although it achieves a median ROC-AUC of 0.64, significantly outperforming both the CNV- and methylation-based models, only slightly below that of the gene expression-based model. This indicates a potential overlap of the prediction-relevant information present in these two data types.

In order to disentangle such potential redundancies in the data and more accurately quantify the individual contributions of data types to the predictive performance of the integrated models, ablation studies are performed. Fig 5 demonstrates the resulting relative mean loss of performance for every second-step algorithm separately: 41 ablation models per second-step algorithm are trained and tested across ten folds and their respective performance is evaluated via ROC-AUC and accuracy for the best cut-off; in the case of the Naïve Bayes classifier, only the latter metric is computed and used for plotting. All of the remaining algorithms are evaluated by ROC-AUC. For each algorithm, the resulting performance metrics of any ablation model are averaged across all folds and then normalized by dividing by the mean performance metric of the corresponding complete two-step model. Consequently, values close to 1 indicate that the removal of a certain set of input data types on average does not affect any particular reduced model, while values smaller than 1 hint at a loss of performance due to information being discarded that cannot be compensated for by the remaining data types.

**Fig. 5.**
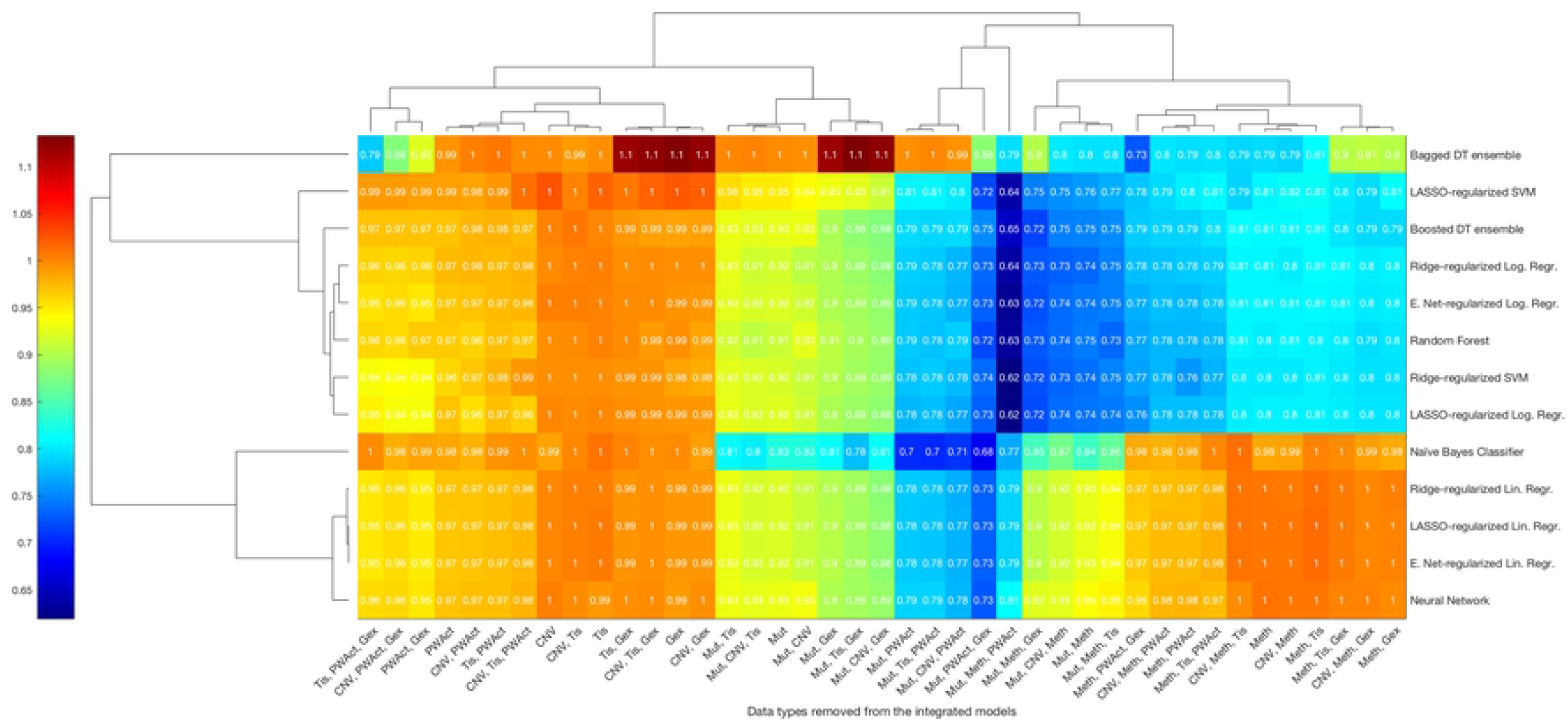
Ablation study results for the compound Nutlin-3a. Heatmap of the mean relative change in performance upon removing combinations of first-step model outputs as input features for the integrated second-step models for Nutlin-3a. Rows correspond to the fitting algorithms used to construct a model, while columns denote the single data type – or combinations thereof – whose corresponding first-step model outputs are removed as inputs. Colours indicate averaged test ROC-AUCs – or accuracy scores in the case of the Naïve Bayes classifier – of ablation models normalized by the respective value of the full model utilizing all available inputs. Values close to 1 indicate that no noticeable change in performance occurred, while scores larger than 1 denote improved predictivity and those smaller than 1 mark a loss in predictivity.

As visible in Fig 5, the removal of CNV, tissue descriptor and gene expression data fails to affect the performance of the integrated models, irrespective of the fitting algorithm applied. Regarding the effect of hypermethylation data, the second-step algorithms are split into two categories: in the case of regularized linear regression approaches, the Naïve Bayes classifier and the Neural Network, removing them results in no quantifiable loss of predictivity, whereas the remaining algorithms suffer a drastic additive loss of performance of around 20%. A universal drop in model performance across all algorithms can be observed in the absence of pathway activation and mutation data, both individually and simultaneously, with the removal of pathway activation data causing a loss of up to 5%, the removal of somatic mutation data generating a loss of up to 11% and their simultaneous removal resulting in a loss of up to 29% in a majority of algorithms. Over all, the regularized linear regression approaches, the Naïve Bayes classifier and the Neural Network approach exhibit the most extreme drop in performance of 27 – 32% upon removing the combination of somatic mutation, pathway activation and gene expression data, while the remaining algorithms suffer most when the combination of somatic mutation, pathway activation and hypermethylation data is not utilized. A notable exception is the bagged decision tree ensemble, which exhibits its lowest predictive performance when hypermethylation, pathway activation and gene expression data are removed from the fitting process.

These findings relate to known pharmacological properties of the compound, most notably its mode of action: *TP53* mutations are identified as statistically significant drivers of resistance in all folds and consequently, the somatic mutation data yield not only the best-performing first-step models, but are also assigned high weights and importance scores by the second-step fitting algorithms. The eleven pathways featured in the PROGENy-supplied pathway activation data include the *TP53* signalling pathway, which is consistently ranked the most significant by the linear regression algorithm applied in the respective first-step model with averaged p-values of around 10^−19^. Other pathways determined to be significant, albeit to a lesser extent, are the MAPK and the *EGFR* pathways. Pathway activation-based first step models constitute the second-best performing first-step models and are assigned high weights or importance scores by a majority of the second-step fitting algorithms. The sole exceptions are the decision tree ensembles and the support vector machines, which favour the mutation- or gene expression-based first-step models exclusively. The aforementioned drastic drop in model performance induced by removing both somatic mutation and pathway activation scores as inputs to the second-step fitting algorithms might implicate that these two data types harbour complementary information pertaining to cellular responsiveness to the compound Nutlin-3a. Considering that the drop in model performance upon removing methylation data does only occur for a subset of algorithms and is not mirrored in the methylation-based model being particularly predictive on its own or even assigned a notable weight or importance score, we assume the effect to be caused mainly by technicalities related to the fitting algorithms and not any underlying biological or pharmacological rationale. This effect indicates that in specific use cases, the choice of fitting algorithm might affect the predictive performance of the resulting models as well as the observed apparent contributions of data types. Analysing the individual features determined to be significant drivers of responsiveness to Nutlin-3a results in confirming well-known factors such as *TP53* mutations and, to a lesser extent, tissue types such as skin [24] or hematopoietic and lymphoid cells [25, 26], that have been shown to react positively to Nutlin-3a treatment. This case study therefore serves as a proof of concept, demonstrating that the features identified as significant by the algorithm do indeed drive the cellular response to the drug. In addition, the models also yield a set of features that, to the best of our knowledge, have yet to be studied in the context of driving tumour responsiveness to Nutlin-3a specifically and therefore constitute promising potential targets for future studies. These are visualized in Fig 6 and include for instance copy number aberrations in *JAK2*, which have already been linked to tumour progression and chemoresistance [27] as well as to responsiveness of lymphoma to a semi-selective kinase inhibitor [28]. A complete list of the features found to be significantly related to shifts of responsiveness can be found in Tables 1–2 in S5 Appendix.

**Fig. 6.**
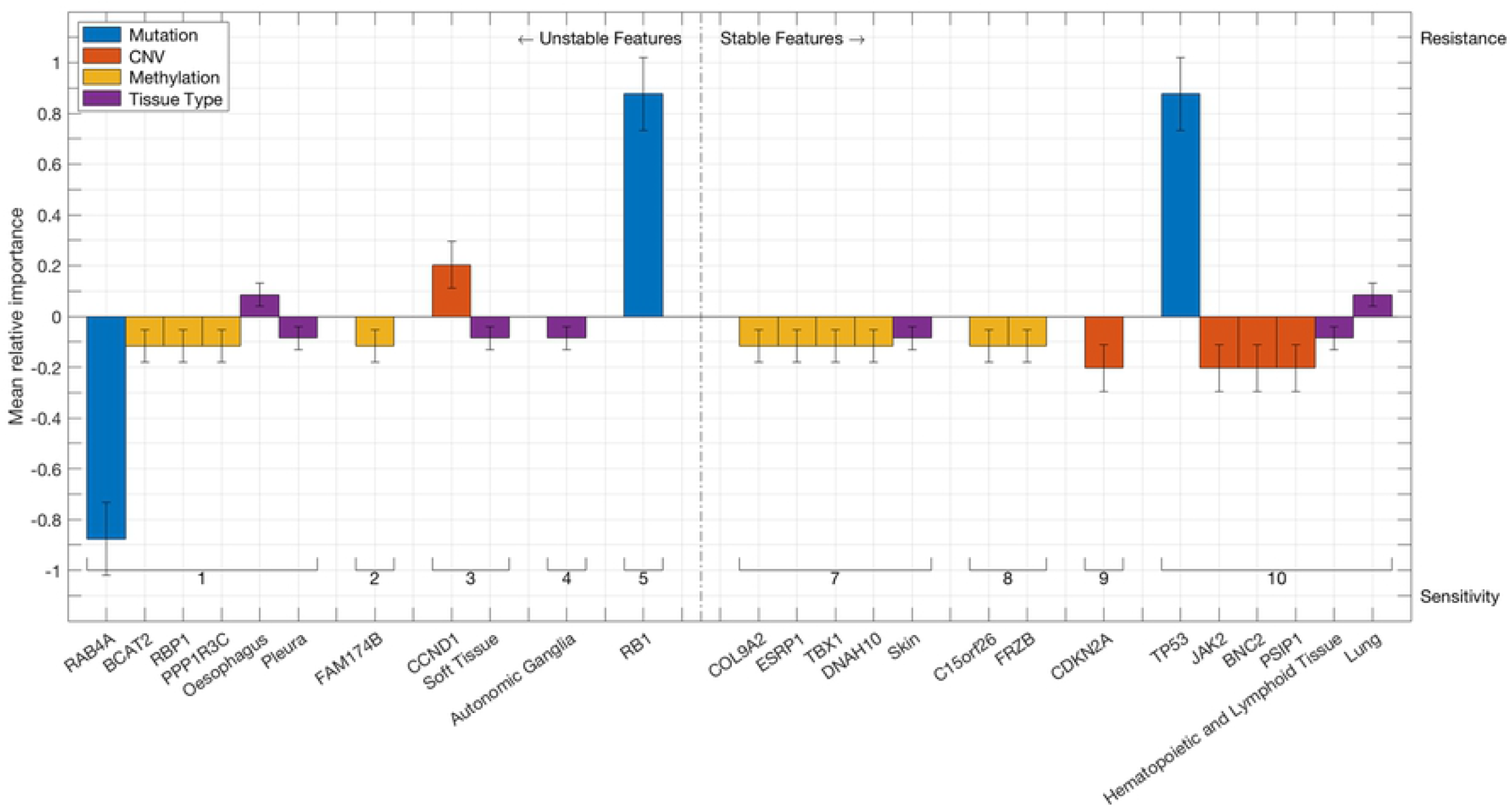
Discrete features linked to responsiveness to Nutlin-3a. Bars represent discrete features that are found to be associated with significant shifts in cellular responsiveness to Nutlin-3a. Colours indicate the data type of the feature, while the absolute bar height corresponds to the mean relative importance of the first-step model based on the respective data type, as calculated by the second-step algorithms. Error bars denote the standard deviation of said importance score. The vertical orientation of any bar indicates whether the associated feature induces sensitivity or resistance to Nutlin-3a, whereas its position on the horizontal axis shows how often it is found in the 10 runs of the cross-validation procedure. Features that are identified at least 7 times out of ten are considered stable.

### Case study 2: Docetaxel

Docetaxel is a cytotoxic chemotherapeutic agent of the taxane family of drugs and is routinely used in the treatment of a wide range of cancers including breast, lung, gastric, colorectal, liver, renal, ovarian, prostate, and head and neck cancers as well as melanoma. Its principal mode of action lies in interfering with the dynamics of microtubule assembly and disassembly, which in turn impedes cell division and promotes apoptosis [29]. In stark contrast to the case of the targeted drug Nutlin-3a, where mutations in the *TP53* gene are well-known to induce resistance, it is not evident from the outset which data types and which corresponding first-step models ought to be expected to perform best in predicting cellular responsiveness to Docetaxel.

Fitting the set of first-step models results in a lack of somatic mutations with a significant link to shifts in responsiveness. CNV- and hypermethylation-based models perform very poorly with averaged test ROC-AUCs rarely exceeding 0.6. Tissue-based models fare moderately better, producing test ROC-AUCs that average 0.74, while gene expression-based and, to a lesser extent, pathway activation-based models, yield models with a high predictive performance with average ROC-AUCs of 0.89 and 0.8, respectively. These findings are visualized in Fig 1 in S6 Appendix. Integrating the first-step model outputs via a Neural Network approach results in models yielding a median ROC-AUC of 0.9 in testing across all folds, slightly outperforming all other second-step fitting algorithms and the gene expression-based first-step model.

As a consequence, the output of the gene expression-based first-step model is consistently determined to constitute the most important and impactful input across all second-step fitting algorithms that allow for an analysis of weights or importance factors, as depicted in Fig 2 in S6 Appendix. Methylation-based and tissue descriptor-based model outputs are assigned a mean relative importance score of 0.24 and 0.2, respectively, while the corresponding scores for CNV-based and pathway activation-based model outputs hover around 0.1. It can be hypothesized that the relatively low importance scores of the comparatively well-performing pathway activation-based and tissue descriptor-based model outputs are due to them containing information that is equally present in gene expression-based model outputs. This redundancy of information content might drive second-step fitting algorithms to assign a high weight only to the gene expression-based input, given that the corresponding first-step model yielded the highest average ROC-AUC. This effect can be observed to be particularly exacerbated in algorithms that have been designed to assign only a small number of non-zero weights to inputs, such as LASSO-regularized regressions.

In order to identify redundancies between data types, the results of the ablation studies for Docetaxel, as visualized in Fig 7, can be evaluated. Since constant first-step models – that is models that fail to find features associated with a shift in responsiveness in a statistically significant way – are not used as inputs to second-step fitting algorithms, the number of ablation models calculated might differ both between distinct drug compounds as well as between different folds for one particular drug. Ablation models that are computed in less than five the folds for any particular drug are excluded in this analysis; as a consequence, the lack of significant somatic mutation features in all but one fold results in only 25 ablation models being calculated with a frequency high enough to warrant further evaluation.

**Fig. 7.**
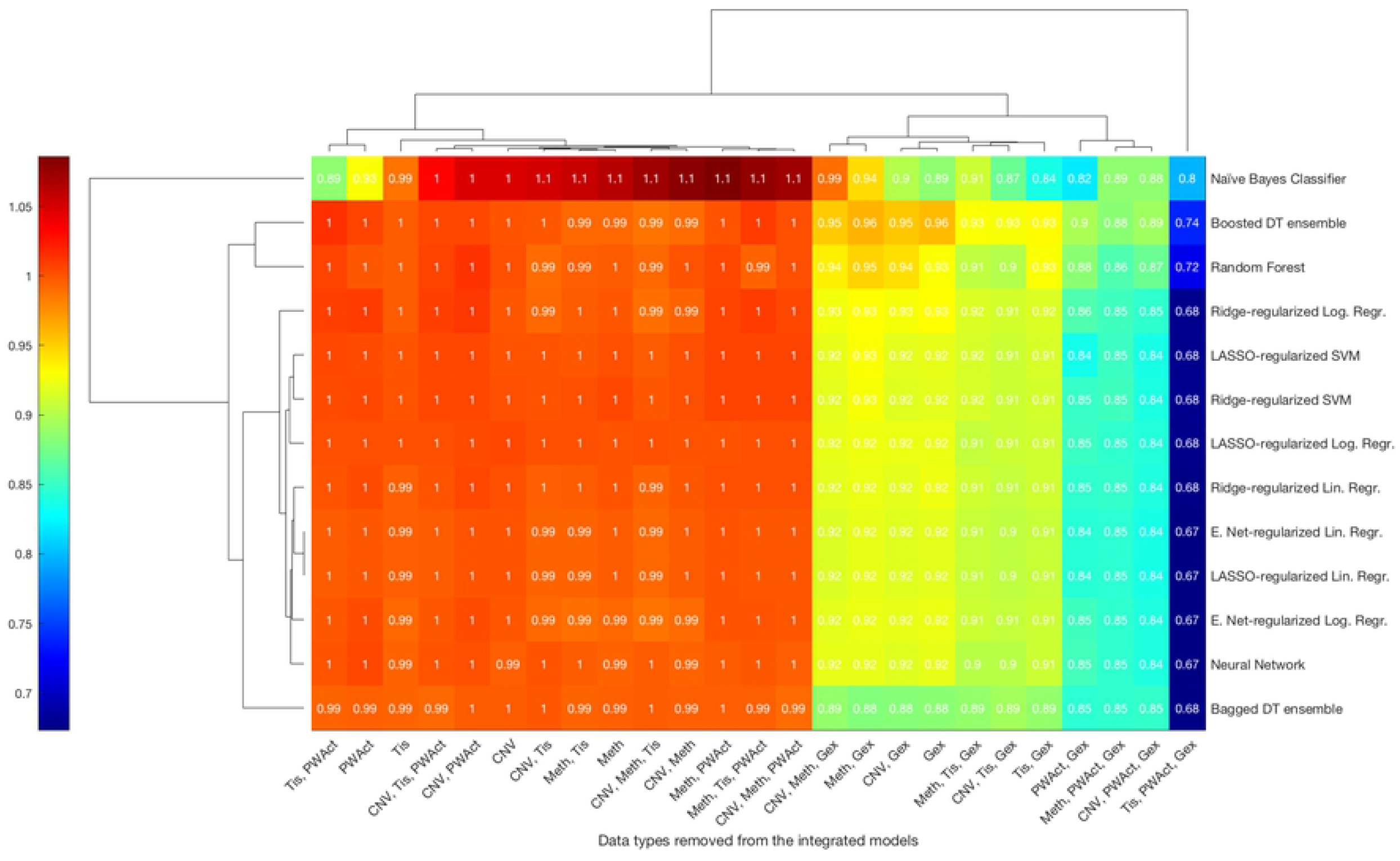
Ablation study results for the compound Docetaxel. Heatmap of the mean relative change in performance upon removing combinations of first-step model outputs as input features for the integrated second-step models for Docetaxel. Rows correspond to the fitting algorithms used to construct a model, while columns denote the single data type – or combinations thereof – whose corresponding first-step model outputs are removed as inputs. Colours indicate averaged test ROC-AUCs – or accuracy scores in the case of the Naïve Bayes classifier – of ablation models normalized by the respective value of the full model utilizing all available inputs. Values close to 1 indicate that no noticeable change in performance occurred, while scores larger than 1 denote improved predictivity and those smaller than 1 mark a loss in predictivity.

The heatmap visualizing the remaining relative average performance of two-step models running on reduced sets of input features shows little difference between distinct second-step fitting algorithms and can easily be divided into two parts. The cluster on the left-hand side of the figure features 14 ablation models that about retain the performance score of the full model with only minor deviations. In contrast, the cluster on the right-hand side consists of eleven models that have been fitted without utilizing the output of the gene-expression based first-step model and that exhibit an increasing decline in performance when viewed from left to right: removing gene expression and any combination of CNV and methylation data results in a drop of performance of about 7 – 8% in the majority of algorithms, while excluding both gene expression and tissue information plus any combination of the aforementioned two additional data types yields a drop of 9 – 10% for a majority of fitting algorithms. In the case of holding out the model outputs based on gene expression, pathway activation and all additional data types but tissue types, a loss of performance of about 14 – 16% can be observed for a majority of algorithms, whereas removing gene expression, pathway activation and tissue descriptor data simultaneously generates a sharp reduction of performance of around 32 – 33% in most algorithms. These findings strongly indicate an overlap among the pieces of information present in these three data types, in particular between the two continuously-valued data types of pathway information and gene expression.

The two-step modelling routine proposed in this paper not only enables the user to analyse and compare the contributions of data types to the overall prediction of the integrated models, but also facilitates the study of the effects of individual features from distinct data types on the distribution of responsiveness across the set of cell lines. Fig 8 depicts the complete set of significant discrete features identified in at least one fold, while a complete list of both discrete and continuous features can be found in Tables 1–2 in S6 Appendix. The set of tissue types found to be linked significantly to a shift in cellular responsiveness to Docetaxel in at least seven out of ten folds includes the upper aerodigestive tract, which is associated with an increase in drug sensitivity and reflects the routine application of Docetaxel in the treatment of head and neck cancers [30]. The list of significant CNVs includes *EGFR*, which is known to constitute a crucial target in cancer therapies in general [31] and that has been shown to drive tumorigenesis in lung cancer when amplifications are present [32]. Relevant and stable hypermethylation events occur, among others, in three members of the *ZNF* family, an extensive set of genes involved in tumorigenesis, cancer progression and metastasis formation [33]. Moreover, the list of crucial sites for hypermethylation events features *WNK4*, a member of the *WNK* signalling pathway that has been linked to cancer progression [34, 35] and is known to interfere with the *TGFB1* pathway. This particular pathway in turn is identified as a significant pathway by the pathway activation models with an average p-value of 10^−4^ and has been found to drive cancerogenesis when misregulated [36]. Additional continuous-valued features include genome-wide expression patterns, namely three principal components calculated on the gene expression data, with averaged p-values of up to 10^−10^.

**Fig. 8.**
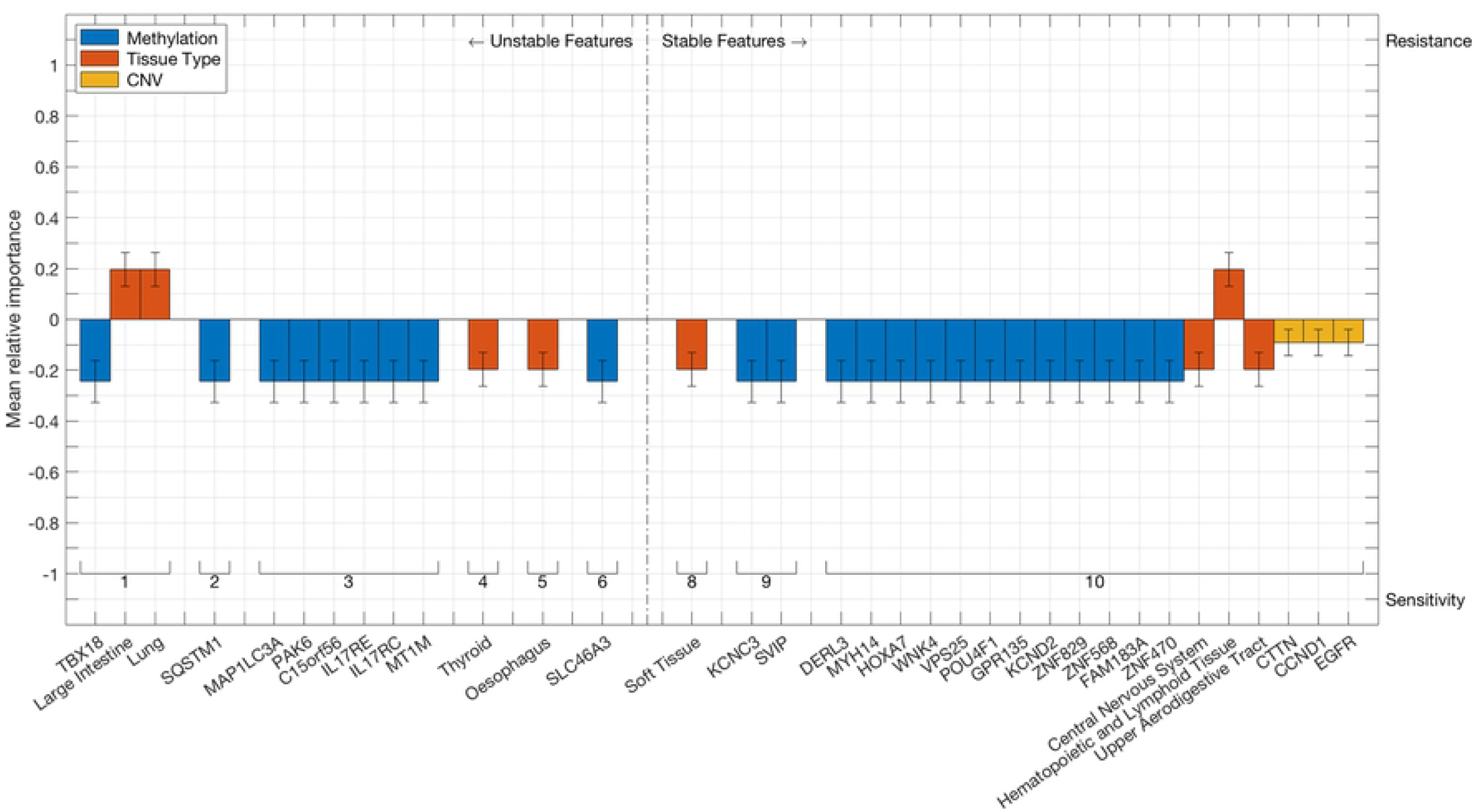
Discrete features linked to responsiveness to Docetaxel. Bars represent discrete features that are found to be associated with significant shifts in cellular responsiveness to Docetaxel. Colours indicate the data type of the feature, while the absolute bar height corresponds to the mean relative importance of the first-step model based on the respective data type, as calculated by the second-step algorithms. Error bars denote the standard deviation of said importance score. The vertical orientation of any bar indicates whether the associated feature induces sensitivity or resistance to Docetaxel, whereas its position on the horizontal axis shows how often it is found in the 10 runs of the cross-validation procedure. Features that are identified at least 7 times out of ten are considered stable.

### Future developments

Currently, the two-step modelling algorithm is specified to run on the GDSC data base as it provides an immense depth of characterization of a wide range of human cancer cell lines that have been tested against a high number of diverse drug compounds. Ideally, in an effort to further prove the reliability of the obtained results and to minimize the chances of them being overfitted to the GDSC data base, one would subsequently run the algorithm on additional large pharmacogenomic data bases which overlap with the GDSC data base in terms of the set of cell lines and drug compounds that are included and the omics data types that are profiled. Unfortunately, to the best of our knowledge, there is currently a lack of publicly available data bases that meet those requirements. The CCLE data set would seemingly constitute the most appropriate candidate; however, systematic analyses [37] have demonstrated that response measurements regarding the small set of shared drug components between both data sets are highly discordant. As a consequence, associations between genomic features and drug response have also been shown to be extremely inconsistent between the two data bases, which effectively renders the CCLE unsuitable to currently perform a comparative study on.

We are looking forward to future publications or modifications of data sets that support a cross-platform study of the stability of results on at least a subset of drug components across a variety of cancer cell lines. However, the applications of the approach presented in this paper do not have to remain limited to pre-clinical human cancer cell line sets. Due the versatility of the underlying principle, the algorithm can easily be adapted to aid research in a broad range of areas, where it is equally imperative to integrate distinct heterogeneous data sources and to identify drivers of pharmacological effects.

## Conclusion

In this paper, we propose a two-step multi-omics modelling approach for the pan-cancer classification of cell lines into responders and non-responders with respect to a wide range of anti-cancer drug compounds. Our algorithm is designed to integrate six distinct data types in a manner that reduces the chance that the process of fitting weights to the input data features is influenced more strongly by structural heterogeneity rather than by the relevant information content. A range of different classification approaches is used for the integration step, which enables users to compare their respective performances. In addition, our algorithm allows for a straightforward in-depth analysis of redundancies between the pieces of information present in the distinct data types and of individual features that shift responsiveness. As a consequence, it produces more interpretable models that not only show a predictive performance that is comparable to the gold standard, but additionally yield valuable biological insights into driving mechanisms and factors. The case studies presented in this paper underscore that our approach succeeds both in correctly identifying established driving features of drugs with a well-understood mechanism of action as well as in finding a set of as of yet unrelated features that constitute suitable candidates for future studies. Comparing our results on the GDSC data set with that of comparable studies implies that our ansatz might be particularly well-suited to be applied to a particular set of targeted drug compounds. Currently, the MATLAB routine is run on the GDSC data base, but the design and implementation of the model is easily generalizable and can be modified and applied to a range of data bases and classification problems.

## Availability of data and materials

The preprocessed datasets analysed during the current study as well as the MATLAB source code of the two-step modelling algorithm are available to be downloaded from the official github repository of the Joint Research Center for Computational Biomedicine, https://github.com/JRC-COMBINE/two-step-modelling [39]. It is licensed under the GNU General Public License v3.0.

## Acknowledgments

Violin plots were generated using a modification of a MATLAB routine developed and made publicly available by H. Hoffman [38].

## Supporting information

**S1 Fig. Extended modelling workflow** Comprehensive figure of the complete modelling workflow..

**S1 Table. Model parameter details** Table detailing the settings of model parameters, as they differ from the default settings provided by MATLAB.

**S1 Appendix. Overview over the model results.** Visualizations of the distributions of model performances; drug compounds are sorted according to their target mechanism.

**S2 Appendix. Overfitting effects for particular algorithms.** Visualization of the effects of low sample numbers on models fitted via Boosted Decision Tree Ensembles and a depiction of overfitting effects as a function of the model training performance.

**S3 Appendix Impact of drug class and algorithm on model performance** Heatmaps visualizing the significance of differences between the model performances on distinct drug classes as well as between model performances of models fitted via distinct algorithms.

**S4 Appendix. Comprehensive comparison of model performances to one-step multi-omics models.** Drug class-specific enrichment p-values of drug compounds on which the two-step modelling approach outperforms the one-step models constructed by Jang et al.; overview of drug compounds where one-step models were found to be more predictive and a comparison of overall distributions of performance.

**S5 Appendix. Discussion of results pertaining to Nutlin-3a** List of discrete and continuous features associated with shifts in sensitivity to Nutlin-3a; visualization of the predictive performances of first-step models and their relative importance, as quantified by second-step algorithms.

**S6 Appendix. Discussion of results pertaining to Docetaxel** List of discrete and continuous features associated with shifts in sensitivity to Docetaxel; visualization of the predictive performances of first-step models and their relative importance, as quantified by second-step algorithms.

